# Ecological tradeoffs lead to complex evolutionary trajectories and sustained diversity on dynamic fitness landscapes

**DOI:** 10.1101/2023.10.11.561986

**Authors:** André Amado, Claudia Bank

## Abstract

The course and outcome of evolution are critically determined by the fitness landscape, which maps genotype to fitness. Most theory has considered static fitness landscapes or fitness landscapes that fluctuate according to abiotic environmental changes. In the presence of biotic interactions between coexisting genotypes, the fitness landscape becomes dynamic and frequency-dependent.

Here, we introduce a fitness landscape model that incorporates ecological interactions between individuals in a population. In the model, fitness is determined by individuals competing for resources according to a set of traits they possess. An individual’s genotype determines the trait values through a Rough Mount Fuji fitness landscape model, allowing for tunable epistasis (i.e., non-additive gene interaction) and trait correlations (i.e., whether there are tradeoffs or synergies in the ability to use resources). Focusing on the effects of epistasis and trait correlations, we quantify the resulting eco-evolutionary dynamics under simulated Wright-Fisher dynamics (i.e., including genetic drift, mutation, and selection under the assumption of a constant population size) on the dynamics fitness landscape in comparison with a similar, static, fitness landscape model without ecological interactions.

Whereas the non-ecological model ultimately leads to the maintenance of one main genotype in the population, evolution in the ecological model can lead to the long-term coexistence of several genotypes at intermediate frequencies across much of the parameter range. Including ecological interactions increases steady-state diversity whenever the trait correlations are not too strong. However, strong epistasis can hinder coexistence, and additive genotype-phenotype maps yield the highest haplotype diversity at the steady state. Interestingly, we frequently observe long-term coexistence also in the absence of induced trade-offs in the ability to consume resources.

In summary, our simulation study presents a new dynamic fitness landscape model that highlights the complex eco-evolutionary consequences of a (finite) genotype-phenotype-fitness map in the presence of biotic interactions.

## 1 Introduction

The concept of fitness landscapes was introduced almost a century ago by Wright [1]. Initially regarded as a metaphor, fitness landscapes sustained long theoretical developments and recently became empirically measurable in the lab [reviewed in 2, 3, 4]. Fitness landscapes promise to be a relevant tool for predicting the evolution of organisms, especially pathogens, whose rapid evolution presents a significant societal challenge, e.g., through the rise and maintenance of resistance to antimicrobial treatments or the emergence of new virus strains [reviewed in 5, 6]. However, the complexity of the fitness landscape of real organisms is gigantic, and even the largest experimental studies are limited to picking a set of genotypes of interest, which are then tested in laboratory conditions [reviewed in 2, 4]. On the theoretical side, there is a need to develop new fitness landscape models that can capture the modular, irregular, and multilayered nature of the true genotype-fitness map and its sensitivity to environmental change [4, 7].

Most existing fitness landscape models assume a static fitness landscape, i.e., that the fitness relationship between genotypes remains constant over time [e.g. 1, 8, 9, 10, 11]. However, fluctuations of the abiotic environment and biotic interactions likely affect the fitness landscape not only regarding the height of its peaks but also regarding their location and the epistatic relationships between genotypes. Indeed, there is increasing evidence that the shape of fitness landscapes can greatly depend on the abiotic environment (e.g., 12, 13, 14, 15, reviewed in 4). The most intuitive way of accommodating abiotic environmental change into fitness landscape modeling is using Fisher’s Geometric model, in which a changed environment can be represented by a changed location of the phenotypic optimum [16]. This extension of the model has inspired several promising works, including evolutionary theory and fitting of experimental data [e.g. 17, 18]. However, the connection between genotype and fitness under Fisher’s Geometric model is not straightforward [19, 20]. Other theoretical studies of fitness landscapes in fluctuating environments, also termed fitness seascapes [7], have relied on genotype-fitness models, the parameters of which were tuned to create temporal, often cyclical, changes in the fitness landscape and to study the resulting evolutionary consequences [e.g. 21, 22, 23], which can be complex [24].

Whereas biotic environmental change has been increasingly incorporated into fitness landscape theory, the influence of biotic interactions on the shape of the fitness landscape has been largely neglected across the fitness landscape research community. Decades ago, Kauffman and Johnsen [25] proposed an elegant extension of their NK fitness landscape model (named after the two main parameters of the model; [11]) to incorporate species interactions, which showed that the presence of species interaction could greatly affect evolutionary dynamics. However, this model (named NKC model for the additional component of species interaction), although highly cited, has not been developed much after the original publication.

Conversely to their absence from the fitness landscape literature, biotic interactions and their consequences play a crucial role in theoretical ecology. However, whereas evolutionary change *per se* is a regular feature of theoretical models in ecology (e.g., in the context of adaptive dynamics approaches e.g. 26), the role of the genetic architecture and genetic constraints remains largely unexplored, with the effects of the number of loci contributing to a trait and the role of epistasis mostly unknown [27]. Some work has focused on one of these aspects in the context of evolutionary rescue [28, 29, 30, 31] or predator-prey models [32, 33, 34], but we are not aware of any work that has incorporated complex fitness landscapes.

Here, we present and analyze a new dynamic eco-evolutionary fitness landscape model, which allows for tunable ecological trade-offs and epistasis. We consider a genotype-phenotype map generated by a modified version of the Rough Mount Fuji fitness landscape model [8]. This model allows us to introduce statistical correlations between traits and tune the degree of epistasis at the trait level. The trait correlation parameter determines whether there are tradeoffs or synergies between traits. The trait values are then mapped to fitness via a resource competition model, which provides a simple means to include frequency-dependent interactions between genotypes. Using simulations under a Wright-Fisher population-genetic model with selection, mutation, and a constant population size, we quantify the conditions for long-term coexistence of multiple intermediate-frequency genotypes. Our dynamic eco-evolutionary model features long-term co-existence of multiple genotypes. As expected, stronger tradeoffs between traits lead to a larger diversity of genotypes in the population at the steady state. However, also in the absence of induced trait correlations, diversity is boosted by ecological interactions, and multiple genotypes can coexist at the steady state. Although steady-state diversity generally decreases with stronger epistasis, phenotypic divergence is maximized for an intermediate strength of epistasis. Interestingly, evolutionary end states were similarly repeatable in the ecological and the non-ecological models, except in the simultaneous presence of epistasis and negative trait correlations. In summary, our model provides an intuitive way of linking genetic and ecological interactions, and our results highlight the importance of considering the role of genetic constraints in the maintenance of long-term diversity.

## 2 Model and Methods

### 2.1 Definition of the dynamic genotype-phenotype-fitness landscape

We model a genotype-phenotype-fitness landscape in which an individual’s genotype determines its carrier’s ability to consume two (or more) resources; we denote this as the phenotype or trait combination. The phenotype determines the fitness of the individual in the current population according to a consumer-resource model.

To model the genotype-phenotype level, we use a multiplicative version of the probabilistic Rough Mount Fuji fitness landscape model [8] to assign phenotypes to genotypes. Mathematically, the phenotype of of an individual with genotype 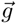 with *L* diallelic loci is

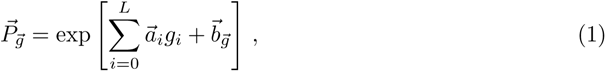

where the additive 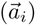 and epistatic 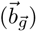 components are drawn from multivariate normal distributions with covariance matrices Σ*_a_* and Σ*_e_*, respectively. The presented definition results in universal pleiotropy, i.e., every mutation affects all traits. Figure 1 provides three examples of possible genotype-phenotype maps illustrating different trait correlations.

**Figure 1:**
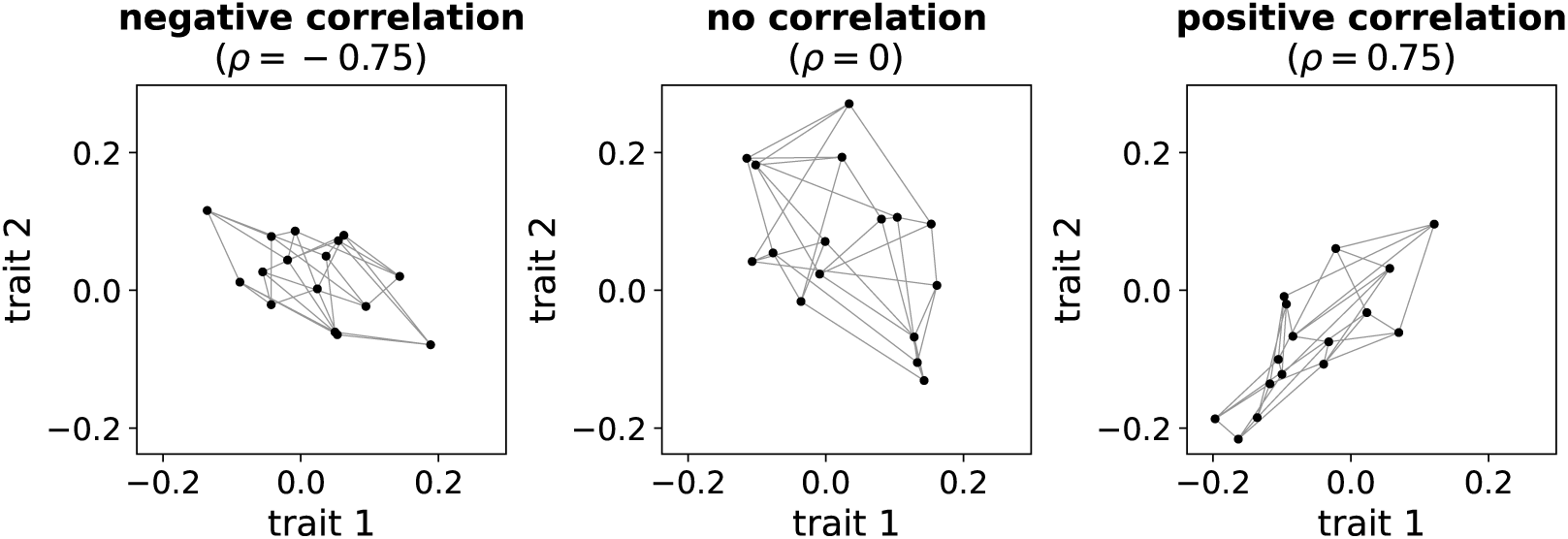
Examples of probabilistic genotype-phenotype maps with different correlations between traits. We denote the map of a genotype to a set of traits (here, a pair) as the genotype-phenotype map. Each dot represents a genotype, and lines connect between single-step mutational neighbors. The parameters are as follows: number of loci *L* = 4, standard deviation of fitness effects *σ* = 0.1, strength of epistasis *e* = 1*/ √*2 (i.e., equal additive and epistatic contribution to the trait variance).

We assume that the phenotype represents the uptake rate of an individual for a set of resources and model the fitness function using a consumer-resource model previously invoked in **(author?)** [35]. Our model considers that individuals compete for a pool of existing nonessential resources; the population density is kept constant over time (see Section Simulation parameters).

In a population of size *N* , an individual with genotype *i* has fitness

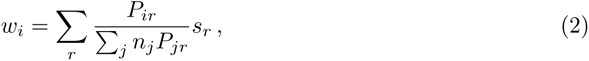

where *P_ir_*stands for the trait value of an individual with genotype *i* for resource *r*, *n_j_* is the number of individuals with genotype *j* in the population and *s_r_* is the amount of resource *r*. The mean fitness is thus given by

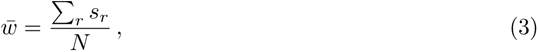

the total amount of resources divided by the population size. Consequently, the relative fitness is

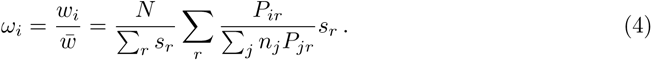

Rewriting this expression in terms of the total amount of resources 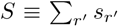 and the mean value of P over the population 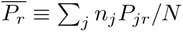 for resource r, the relative fittness simplifies to

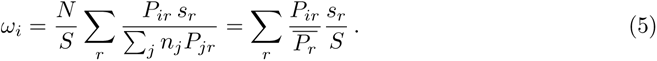

Given any current distribution of genotypes, we combine the genotype-phenotype and the phenotype-fitness map to build a genotype-fitness landscape. This fitness landscape depends on the frequency distribution of all genotypes. Therefore, although the phenotype associated with a particular genotype remains the same over time (i.e., there is no phenotypic plasticity), the fitness landscape that an evolving population perceives changes over time (see Supplementary Video 1).

To compare results with and without ecological interactions, we introduce a non-ecological model with a static fitness landscape. Here, we determine fitness as the sum of the trait values of a genotype instead of using the weighted average resulting from the competition for resources. Supplementary Figure S1 shows that the statistical properties of the fitness landscapes [mean fitness effect, variance of fitness effects, and gamma epistasis statistics (a measure of epistasis based on the correlation of fitness effects across genetic backgrounds; see [36] for a detailed discussion of this statistic)] of the ecological and the non-ecological models are similar across most of the parameter range.

### 2.2 Evolution of the population on the fitness landscape

We consider haploid populations of constant size *N* , which evolve on the dynamic fitness landscapes and their static counterparts. We initiate the simulations with a monomorphic population occupying a genotype chosen randomly with uniform probability. Individuals undergo recurrent mutation with a probability of *µ* independently per locus per generation, after which the population undergoes a genotype frequency update according to a Wright-Fisher model with selection. Here, a new population completely replaces the previous one, with the new individuals being offspring of the old ones with a probability proportional to their fitness, and keeping the population size *N* constant. This update introduces selection and random genetic drift to the evolutionary process.

### 2.3 Simulation parameters

We simulated the evolution of populations on the dynamic and static fitness landscapes for a range of parameters. We considered fitness landscapes with *L* = 10 loci for most results in the paper and provided the same results for lower-dimensional fitness landscapes with *L* = 5 in the Supplementary Information. We used a fixed population size of *N* = 1000 individuals and a mutation rate of *µ* = 0.001. We considered two resources throughout the paper and fixed the resource supply at *s_r_* = 1 for both resources. This situation is analogous to a chemostat in an experimental context, where resource availability is kept constant over time.

The mean effect of mutations on the phenotype was set to zero, and their variance *σ*^2^ was kept constant (*σ* = 0.1). The variance of effects on phenotype was then distributed between additive and epistatic components of the Rough Mount Fuji model. We introduced a single parameter *e* with a range of 0 to 1 that summarizes the strength of epistasis in the genotype-phenotype landscapes. Here, the epistatic variance *σ_e_*^2^ is defined as *σ*^2^*e*^2^*/*2, and the additive variance *σ_a_*^2^ is *σ*^2^(1 *− e*^2^). Since the variance of the effects on the trait is given by

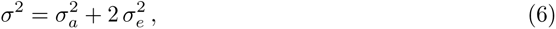

the above definitions allow us to maintain a constant effect on the phenotype while varying the strength of epistasis *e*. In other words, increasing *e* converts additive variance into epistatic variance while keeping the total invariant.

The parameter *ρ* determines the correlation between traits of the two genotype-trait landscapes. Large positive *ρ* represents a highly correlated pair of genotype-trait landscapes, large negative *ρ* an anticorrelated pair of genotype-trait landscapes, whereas *ρ* close to zero generates uncorrelated genotype-trait landscapes.

Written in terms of the above parameters, the covariance matrices Σ*_a_* and Σ*_e_* take the forms

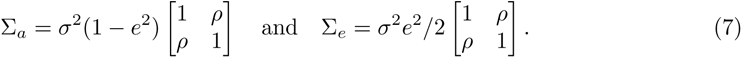

We analyzed simulations from a range of *e* and *ρ* values to disentangle the effects of epistasis and trait correlation on long-term diversity.

### 2.4 Characterization of the final state of the population

Due to their stochasticity and the occurrence of recurrent bidirectional mutation, our simulations can by design never reach a deterministic equilibrium. We therefore defined a criterion for successful convergence to a (semi-)stable state in order to quantify long-term diversity and the repeatability of the evolutionary process. We characterized the final state by the set of genotypes it contains. In order to qualify for being considered in the set of genotypes of a final state, a genotype should have a nonzero frequency and an above-average relative fitness over a specified diagnostic period (here, 500 generations; this number was determined through manual exploration). We excluded any genotypes that fulfilled this criterion for less than 10% of the diagnostic period from the final state because they tended to represent recurrent single-step mutational neighbors of stable high-fitness genotypes. Once the set of genotypes defined by these criteria remained constant for 500 consecutive generations, we determined that the system had reached its final state. We used a burn-in period of 15000 generations to guarantee exploration of the genotype space in every simulation. All simulations reached their final state within 100000 generations.

### 2.5 Diversity measures

To quantify the genetic diversity in the population throughout the simulation and at the final state, we computed the haplotype diversity. Note that since we consider haploid individuals throughout the paper, haplotypes and genotypes are equivalent. We defined haplotype diversity as

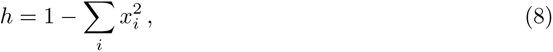

where *i* indicates the genotype and *x_i_*is the fraction of individuals carrying genotype *i*. This quantity represents the probability that two individuals randomly selected from the population carry different genotypes.

We also quantified the diversity of the population at the phenotypic level. For that, we used the mean phenotypic distance, i.e., the mean Euclidean distance between the phenotypes of two randomly chosen individuals. We defined the mean phenotypic distance as

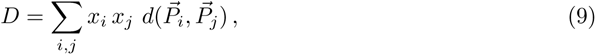

where *i* and *j* are genotype indices, *x_i_* is the frequency of genotype *i* in the population, *d*(*·, ·*) is the Euclidean distance, and 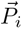 the phenotype corresponding to genotype *i*.

### Code Availability

All annotated simulation code to reproduce the study will be made available on Zenodo upon publication.

## 3 Results

We follow the evolution of a haploid population of *N* individuals that is initially monomorphic for a random genotype. Over time, individuals mutate and compete according to their relative fitness. As mutation generates new genotypes, the population experiences an initial diversification. This initial spike in diversity happens at the genotypic and phenotypic levels and is similar for both eco-evolutionary and non-ecological models. Subsequently, evolution proceeds differently for each model. An example simulation run is depicted in Fig. 2, accompanied by a video of an evolving fitness landscape provided in the Supplementary Information.

**Figure 2:**
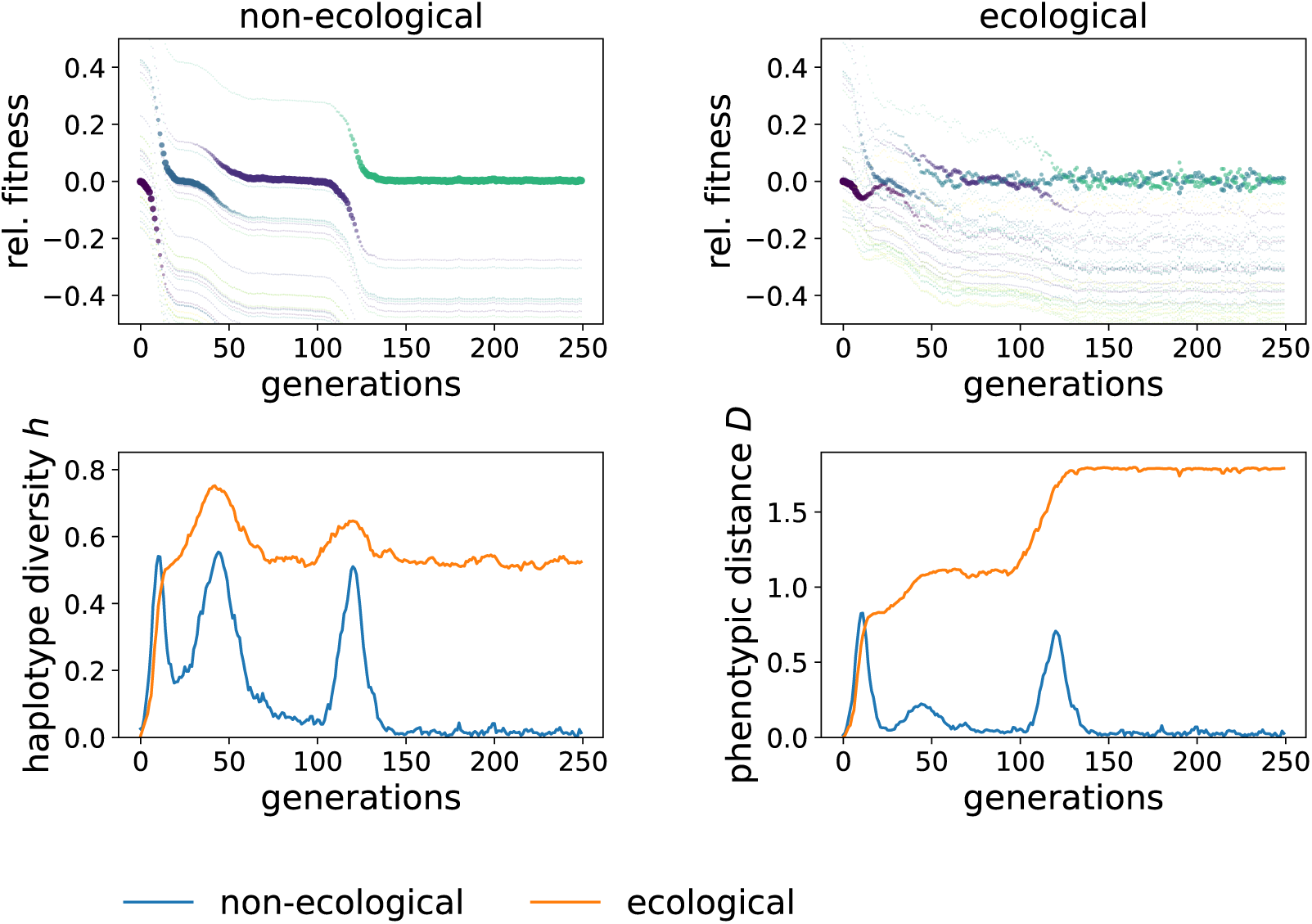
Evolutionary dynamics differ greatly between the ecological and the non-ecological models. The top row depicts the relative fitness as a function of time for all genotypes in the fitness landscape. The dot radius is proportional to the frequency of the respective genotype in the population. The left panel shows the result for the non-ecological model and the right panel for the model with ecological interactions. The bottom row displays the diversity over time for the same simulation runs, with the non-ecological model shown in blue and the ecological model in orange. The left graph depicts the haplotype diversity, and the right one the phenotypic distance. The strength of epistasis *e* is chosen so that the additive and epistatic components have an equal contribution to the effects on phenotype (*e* = 1*/ √*2). The remaining parameters are the number of loci *L* = 5, the population size *N* = 1000, the trait correlation *ρ* = *−*0.9, the mutation rate per locus *µ* = 0.001, the resource amount *s_r_* = 1, and the variance of fitness effects *σ* = 0.1.

### 3.1 Evolution on the dynamic fitness landscape features fuzzy sweeps of multiple genotypes without loss of diversity

In the non-ecological model, we observe series of selective sweeps, where fitter genotypes sequentially invade and replace the previous genotypes until evolution becomes stalled at a local or the global fitness peak. There are temporary spikes of diversity during the selective sweeps, but the fixation of new genotypes erases this temporary diversity. Once a peak is reached, both genotypic and phenotypic diversity remain stable at low values.

In the ecological model, the situation is more complex. Here, several genotypes compete simultaneously. In many cases, the population reaches a polymorphic equilibrium with two or more coexisting genotypes at intermediate frequencies. In this case, the initial diversification introduces new resource acquisition strategies that modify the competition for resources and thus alters the fitness landscape experienced by the population. There is a transient period during which local maxima arise and vanish as the evolving population renders them unstable. As seen in Fig. 2, selective sweeps are less clear and can involve several genotypes simultaneously (e.g., around generation 50 in the example shown in Fig. 2). During these sweeps, haplotype diversity shows less pronounced peaks than in the non-ecological model. Here, the phenotypic distance increases but does not drop after the sweep, in sharp contrast with the non-ecological model where a steep drop follows the temporary increase of diversity during a selective sweep (e.g., around generation 125 in the example shown in Fig. 2). Typically after a few hundred generations, populations evolving under the ecological model reach the final state. Here, a set of intermediate-frequency genotypes resides or fluctuates at a dynamic fitness peak, often preserving a stable coexistence.

At the final state, the fitness rank order of the intermediate-frequency genotypes tends to fluctuate over time because genetic drift and selection during the competition for resources continually alter genotype frequencies and fitness. However, populations sustain these oscillations in a stable fashion for thousands of generations. Supplementary Video 1 shows the evolution of an example fitness landscape over time.

### 3.2 Repeatability of final states is largely unaffected by ecological interactions

Different final states can be reached for the same underlying phenotypic landscape depending on the contingencies of the eco-evolutionary trajectory. Figure 3 (*L* = 10 loci) and Supplementary Fig. S2 (*L* = 5 loci) quantify the repeatability of the final state, i.e., the probability of reaching the same final state in two independent runs. For each fitness landscape, we performed 100 simulations, each starting from a random initial genotype, and averaged over 100 different fitness landscapes for each set of parameters. As expected, repeatability is high for small landscapes (*L* = 5 loci) irrespective of the remaining parameters since the explorable space is small. On larger landscapes (*L* = 10 loci), repeatability is large for additive landscapes but decreases as the underlying phenotypic landscapes become more epistatic. Very epistatic landscapes impede the exploration of the genotype space, and the population often ends up trapped in different local maxima that depend on the specific path taken in each replicate. Interestingly, ecological and non-ecological models show similar repeatability over a large range of parameters; only when epistasis is strong (but not too strong) and trait correlations are negative, the repeatability of the final state is consistently lower in the ecological than in the non-ecological model.

**Figure 3:**
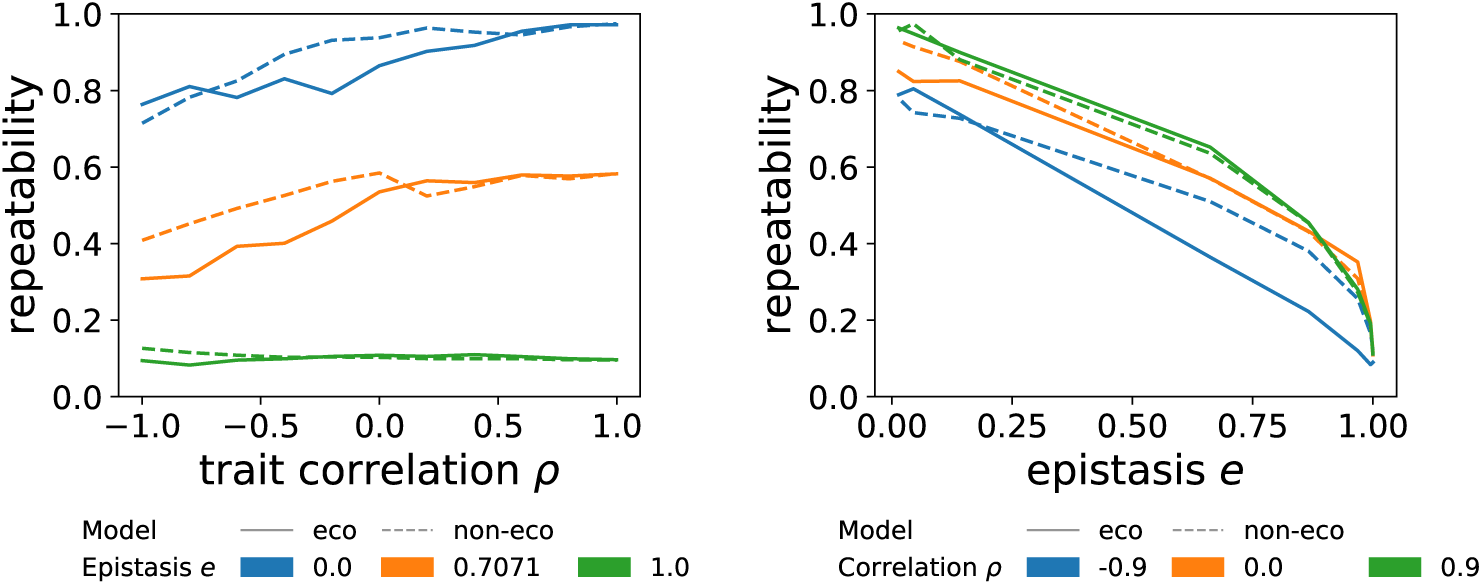
Repeatability of the final states tends to be similar between the ecological and the non-ecological models, unless epistasis is strong (but not too strong) and trait correlations are negative. The left graph shows the repeatability as a function of epistasis *e* under constant trait correlation *ρ* and the right graph as a function of correlation *ρ* for constant epistasis *e*. The repeatibility has been accessed over 100 replicates and averaged over 100 different fitness landscapes with the same parameters. The fixed parameters are the number of loci *L* = 10, the population size *N* = 1000, the mutation rate per locus *µ* = 0.001, the amount of resources *r_i_* = 1, and the variance of fitness effects *σ* = 0.1.

### 3.3 The dynamic fitness landscape facilitates long-term coexistence of multiple phenotypically divergent genotypes

Evolution never ends in our chosen evolutionary modeling approach with recurrent mutation and genetic drift, since genotype frequencies change every generation and mutations are continually (re-)introduced. We therefore quantified a (quasi-)steady state by identifying a set of genotypes that were maintained over a prolonged period of time (here, 500 generations). We further validated that the genotypes in this set were beneficial during at least 50 generations in this diagnostic period to eliminate deleterious genotypes that arise and are maintained at low frequencies solely by mutation. We then computed genetic and phenotypic diversity for the set of stably maintained genotypes (see Materials and Methods) and compared the results to those from the same parameter combinations in the non-ecological model.

Across almost the whole parameter range, the haplotype diversity *h* at the final states was larger for the ecological than the non-ecological model, indicating that coexistence of multiple genotypes with similar fitness became likely. Only when the trait correlation was very large, haplotype diversity was similar between the models. That is expected because, in the limit of large trait correlations, the ecological and non-ecological models become identical. In this limit, *P_ir_* becomes the same for all resources *r*. Therefore, the relative fitness (Eq. 5) simplifies to *ω_i_* = *P_i_/P̄* for both ecological and non-ecological models. Figure 4 shows the haplotype diversity as a function of the strength of epistasis *e* for different values of trait correlation *ρ*. When the trait correlation was very negative (*ρ* = −0.9), haplotype diversity was much larger in the ecological model compared to the non-ecological model. The variance in haplotype diversity was very small for very negative trait correlations, meaning that all genotype-phenotype landscapes produced under the same set of parameters produced similar levels of haplotype diversity. When the traits were uncorrelated (*ρ* = 0), the variance in observed haplotype diversity between genotype-phenotype landscapes produced under the same set of parameters became much wider. Despite the increase in variation, the ecological model on average supported much greater haplotype diversity than the non-ecological model.

**Figure 4:**
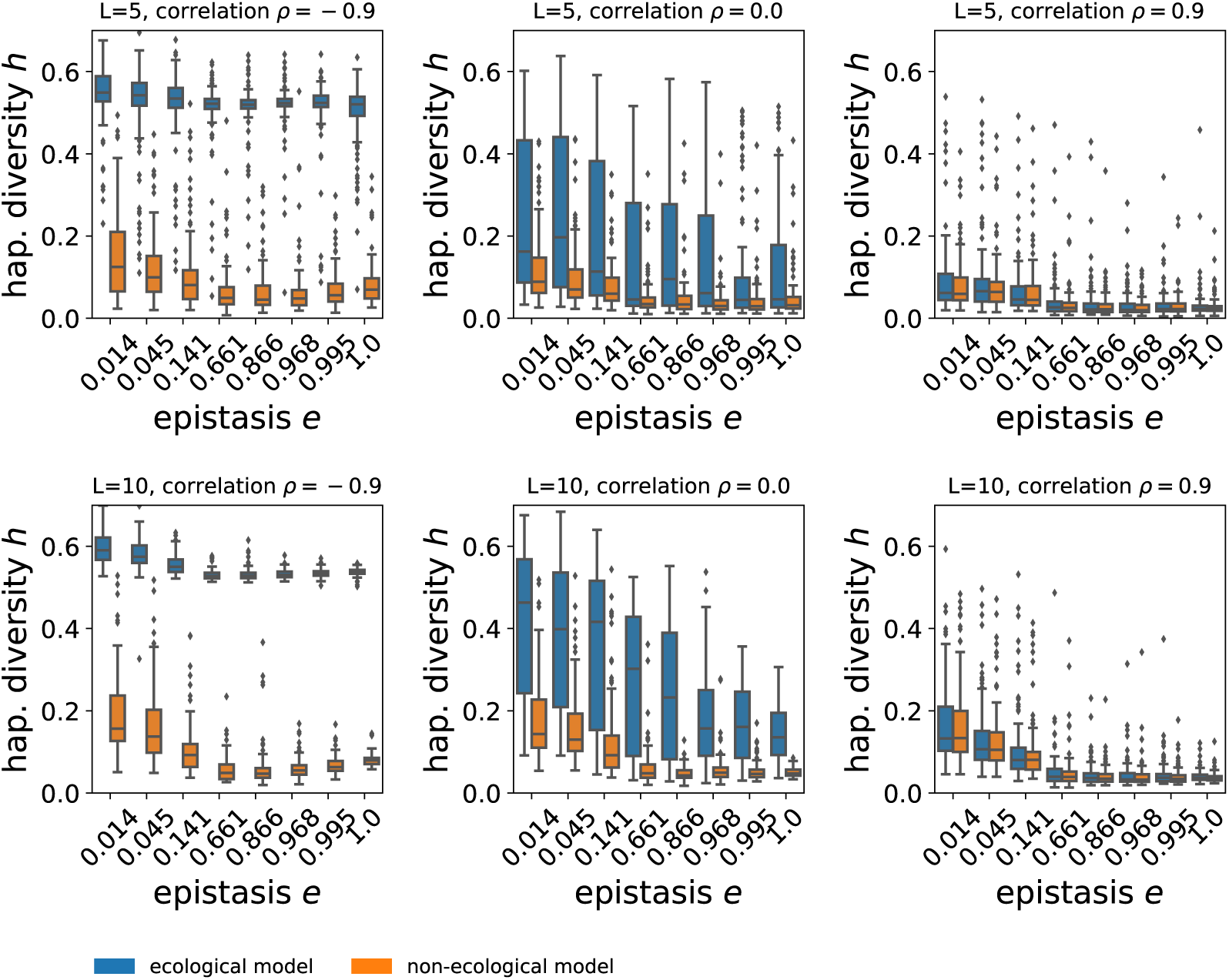
Haplotype diversity tends to decrease with increasing strength of epistasis. The figure shows the haplotype diversity *h* as a function of the strength of epistasis *e* for three values of trait correlation, *ρ* = −0.9, 0, and 0.9. The blue boxes represent the ecological model, and the orange boxes the non-ecological model. The top row shows the results for *L* = 5 loci, whereas the bottom row shows results for *L* = 10 loci. Each point represents data obtained from 100 different fitness landscapes with the same parameters with 50 replicates per landscape. The remaining parameters are the population size *N* = 1000, the mutation rate per locus *µ* = 0.001, the amount of resources *s_r_* = 1, and the variance of fitness effects *σ* = 0.1.

Epistasis tended to affect haplotype diversity negatively (see Supplementary Figure S3). In completely additive fitness landscapes (*e* = 0), both the ecological and the non-ecological models showed the highest haplotype diversity across all values of trait correlations. As epistasis increased, its strength either had no or a negative effect on the haplotype diversity at the final state (see Figure 4). Interestingly, for large negative trait correlations, intermediate epistasis minimizes the haplotype diversity in the non-ecological model. Thus, in the case of large negative interactions, the haplotype diversity gain achieved by the introduction of ecological interactions is maximized for intermediate epistasis (see Figure 4, left panels).

We quantified the mean phenotypic distance *D* at the final state as a measure of functional diversification. Unsurprisingly, the phenotypic distance increased with decreasing trait correlations for any strength of epistasis. This reflects the introduced tradeoffs in the genotype-phenotype landscape which lead to an increasing probability of the establishment of specialist instead of generalist genotypes at the final state. In contrast to the haplotype diversity discussed above, phenotypic distance was maximized for small but non-zero strengths of epistasis in the ecological model (Supplementary Figure 5). Interestingly, for very strong epistasis, the achieved phenotypic distance in the ecological model at the final state was very small across all values of trait correlations, indicating that strong epistasis can hinder ecological diversification even in the presence of strong phenotypic trade-offs.

## 4 Discussion

Although ecology and evolution are tightly intertwined, models of fitness landscapes to date have not provided simple means to incorporate ecological interactions. Here, we developed and analyzed a fitness landscape model that addresses this gap by combining a probabilistic fitness landscape model on the genotype-phenotype level with a consumer-resource ecological model. This resulted in dynamic, frequency-dependent genotype-fitness landscapes that showed complex eco-evolutionary dynamics and featured long-term maintenance of diversity impossible to achieve with static fitness landscapes.

### 4.1 Complex evolutionary dynamics on a dynamic fitness landscape

Even in supposedly simple environments, evolution brings about complex evolutionary dynamics, including ecological diversification [37]. Our model captures such complex evolutionary dynamics by showing that even with just two resources, the population-genetic trajectories and the longterm evolution of a population are greatly changed (Fig. 2) as compared with evolution in a static landscape with the same statistical properties. Our model therefore provides a testing ground for evolutionary biologists to extract how selective sweeps and other population-genetic measures are altered by the addition of ecological interactions.

### 4.2 Maintenance of diversity without trade-offs in a finite genotype space

An important question in ecology is whether multiple species or genotypes can coexist in a given environment. Traditionally, the competitive exclusion principle stated that only *N* species can coexist on *N* resources [38]. However, recent theoretical work has shown that this principle does not generally hold [35, 39, 40]. In particular, when tradeoffs in the ability to consume different resources are imposed, resource-consumer models can sustain an arbitrarily high number of coexisting species [35]. However, these recent models rely on strong assumptions, namely that all organisms have a maximum “enzyme budget” that generates a linear tradeoff between each cell’s capacity to acquire and process different resources.

Our dynamic fitness landscape model differs from the above previous work by incorporating a finite genotype-phenotype-fitness map with tunable epistasis. Interestingly, we find that a large amount of haplotype diversity can be maintained also in the absence of tradeoffs. Unfortunately, it is difficult to compare our results with the requirements of the competitive exclusion principle, since mutation in our model continually (re-)introduces mutations. This leads to a high alpha diversity (i.e., the number of haplotypes present at the final state) both in the ecological and the non-ecological models (see Supplementary Figure S4). However, the mean alpha diversity in the ecological model ranges as high as *>* 15 in the ecological model with 10 loci, which indicates many candidates for coexistence of multiple genotypes. As a crude measure of the part of alpha diversity that corresponds to actual stable genotypes, we could use the difference in the mean number of genotypes present in the ecological versus the non-ecological model (the difference between the graphs in the rows of Supplementary Figure S3). For the parameters that produce the largest differences, we find that there are, on average, 5 extra coexisting genotypes for *L* = 5 and 6 extra coexisting genotypes for *L* = 10. Future work should quantify in which conditions our model results in credible instances in which the competitive exclusion principle was violated. We hypothesize that our finite genotype space that constrains evolutionary paths (both with and without epistasis) should promote the coexistence of multiple genotypes beyond the number of resources.

### 4.3 Epistasis as a hindrance for haplotype diversity but an engine for phenotypic divergence

Although its prevalence in genotype-phenotype or genotype-fitness maps is increasingly accepted, the role of epistasis in the evolutionary process is still debated. We showed that epistasis affects genetic and phenotypic diversity differently in our dynamic fitness landscape model. Whereas an additive genotype-phenotype map maximized haplotype diversity (Supplementary Fig. S3), a small but non-zero strength of epistasis maximized the mean phenotypic distance (Fig. 5). Since we kept the total effect sizes in the genotype-phenotype landscape constant, we hypothesize that single-step mutations may be, on average, more deleterious in fitness landscapes with intermediate epistasis, which could explain the reduced haplotype diversity as compared with an additive fitness landscape. The presence of fewer genotypes in the final state could then lead to a higher mean phenotypic difference between the top specialist genotypes. This hypothesis is supported by the finding that in the non-ecological model, haplotype diversity is minimal for an intermediate strength of epistasis (Fig. 4). These results highlight the importance of including genetic constraints and interactions imposed by epistasis into ecological and evolutionary models.

**Figure 5:**
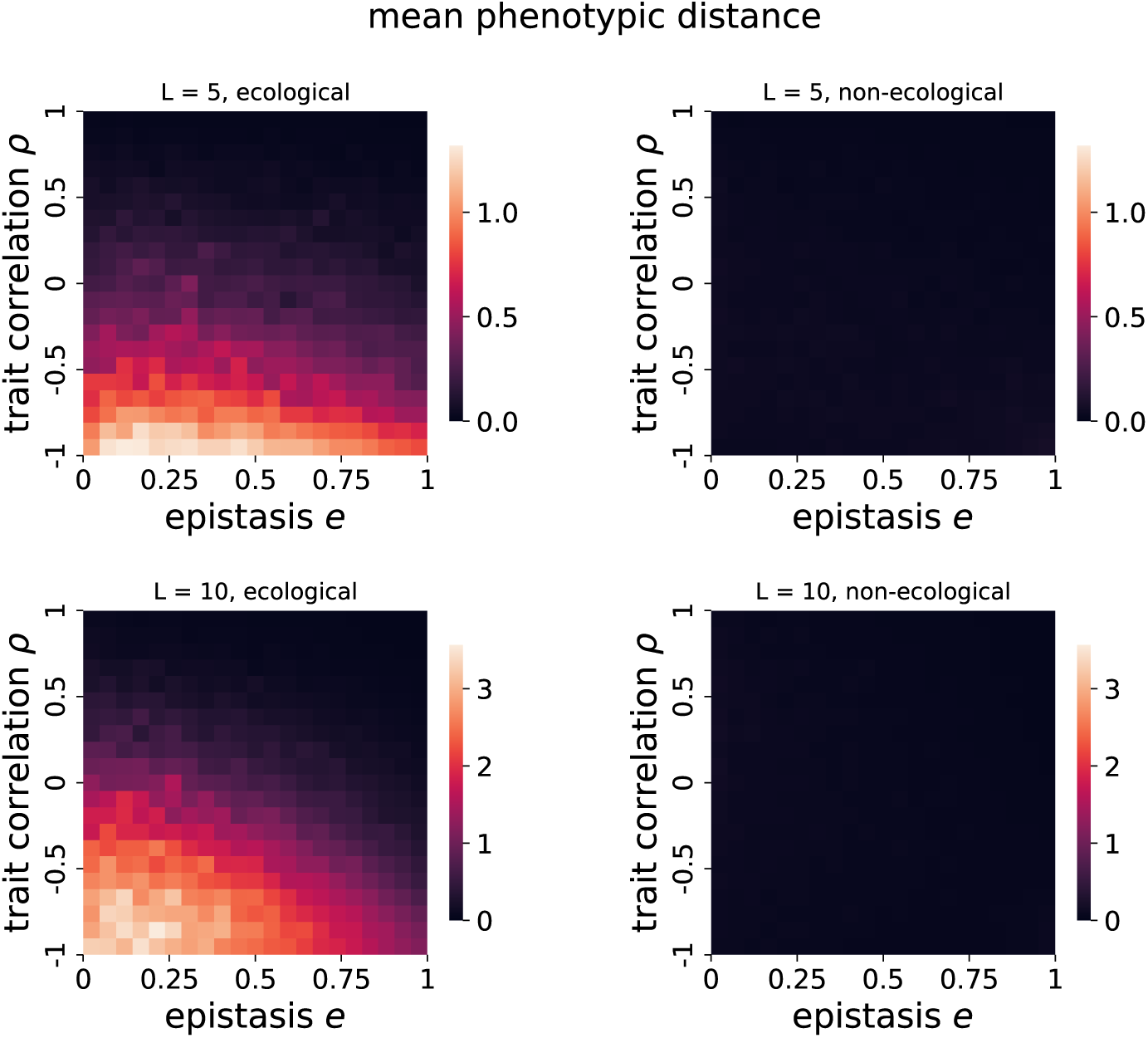
Mean phenotypic distance is maximized for small but positive strengths of epistasis in the ecological model. The graphs show the mean phenotypic distance *D* at the final state. The top row shows the results for *L* = 5 loci and the bottom row for *L* = 10 loci. The left column shows the ecological model and the right column the non-ecological model. Each point represents an average obtained from 100 different fitness landscapes with the same parameters with 20 replicates per landscape. The remaining parameters are the population size *N* = 1000, the mutation rate per locus *µ* = 0.001, the amount of resources *s_r_*= 1, and the variance of fitness effects *σ* = 0.1.

### 4.4 Strengths and limitations of the model

We here presented the prototype of a dynamic genotype-phenotype-fitness model with tunable epistasis and trait correlations. We simulated evolution on the dynamic fitness landscapes using a Wright-Fisher model with selection. This allowed us to simulate evolutionary processes with selection, mutation, and genetic drift, which we could directly compare with evolution on a static fitness landscape with the same statistical and evolutionary features, singling out the effects of incorporating ecological interactions. Our study relies on simulations of a limited set of parameters, which impede the generalization of our results. Moreover, our mutation parameter *µ* = 0.001 was chosen such that it lies in the limit between low and high mutation rates (*Nµ* = 1), potentially generating results that are somewhat specific to this transition regime. Finally, for simplicity, we kept both the population size and the availability of resources constant.

Despite the limitations of our study, we consider our model a good starting point for a theory of dynamic eco-evolutionary fitness landscapes. Under simplifications of the evolutionary dynamics (e.g., with a deterministic approach), it could be possible to derive analytical properties of the model, for example, regarding steady states and their stability. Moreover, the model can be easily modified to feature any kind of genotype-phenotype map (e.g., a Gaussian genotype-phenotype map like in Fisher’s Geometric model, or Michaelis-Menten kinetics to model more mechanistically motivated genotype-phenotype maps). Introducing resource and population dynamics is similarly straightforward, although it will create new statistical challenges and model outcomes. Importantly, our model incorporates tunable parameters of biological interest, such as the genotypic complexity of traits, epistasis, and trait correlations, into the complex framework of a genotype-phenotype-fitness landscape, which can become a testing ground for their effect on eco-evolutionary trajectories in the presence of ecological interactions.

## Acknowledgements

We thank the members of the THEE division for useful discussions throughout the project. This work was supported by funding from ERC Starting Grant 804569 (FIT2GO), SNSF Project Grant 315230/204838/1 (MiCo4Sys), and HFSP Young Investigator Grant RGY0081/2020 to CB.

---

**Figure S1:**
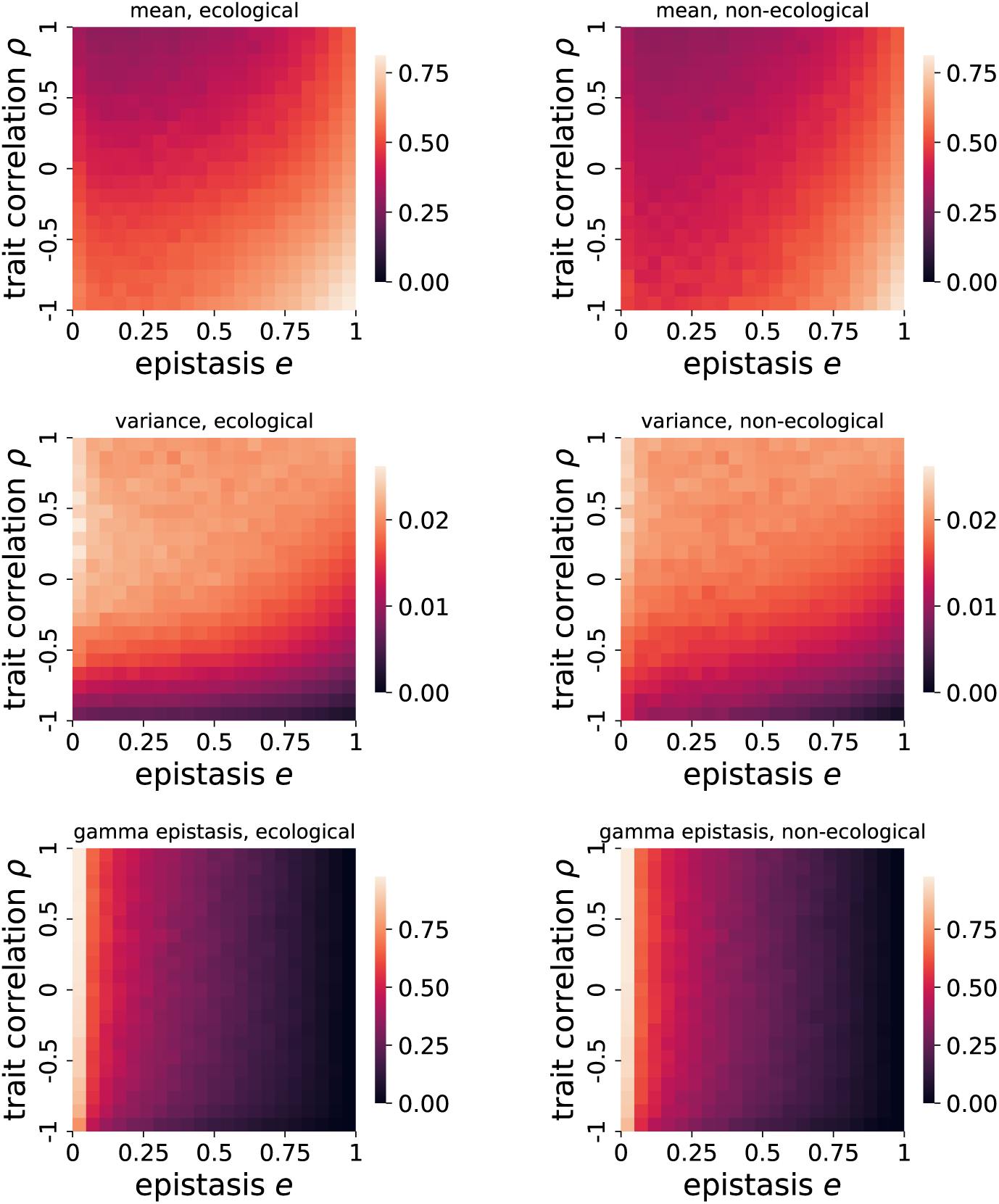
Fitness landscape statistics, averaged over the last 500 generations of each run. The first row shows the mean fitness effect, the second row the variance of the fitness effects, and the third row the gamma statistics (which summarizes the epistasis in the landscape; a value of one indicates no epistasis, whereas a value of 0 indicates strong epistasis). The left column shows the results for the ecological model, and the right column for the non-ecological model. The mean fitness effects show a Pearson correlation of 0.959, a variance of fitness effects of 0.977 and a gamma epistasis of 0.998, indicating a high degree of similarity. Each point represents an average obtained from 100 different fitness landscapes with the same parameters with 20 replicates per landscape.The parameters are as follows: number of loci *L* = 10, population size *N* = 1000, mutation rate per locus *µ* = 0.001, amount of resources *r_i_* = 1, the variance of fitness effects *σ* = 0.1.

**Figure S2:**
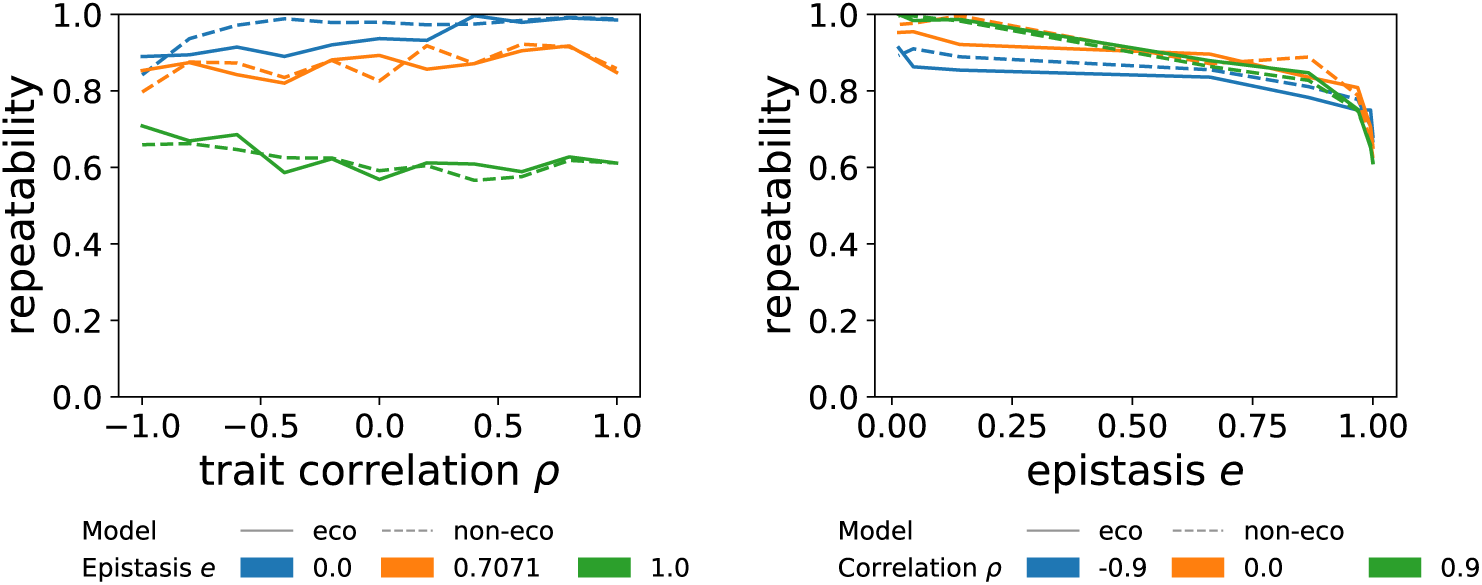
Repeatability of the final states. The left graph shows the repeatability as a function of epistasis *e* under constant trait correlation *ρ* and the right graph as a function of correlation *ρ* for constant epistasis *e*. The repeatibility has been accessed over 100 replicates and averaged over 100 different fitness landscapes with the same parameters. The parameters are *L* = 5 loci, population size *N* = 1000, mutation rate per locus *µ* = 0.001, and amount of resources *r_i_* = 1.

**Figure S3:**
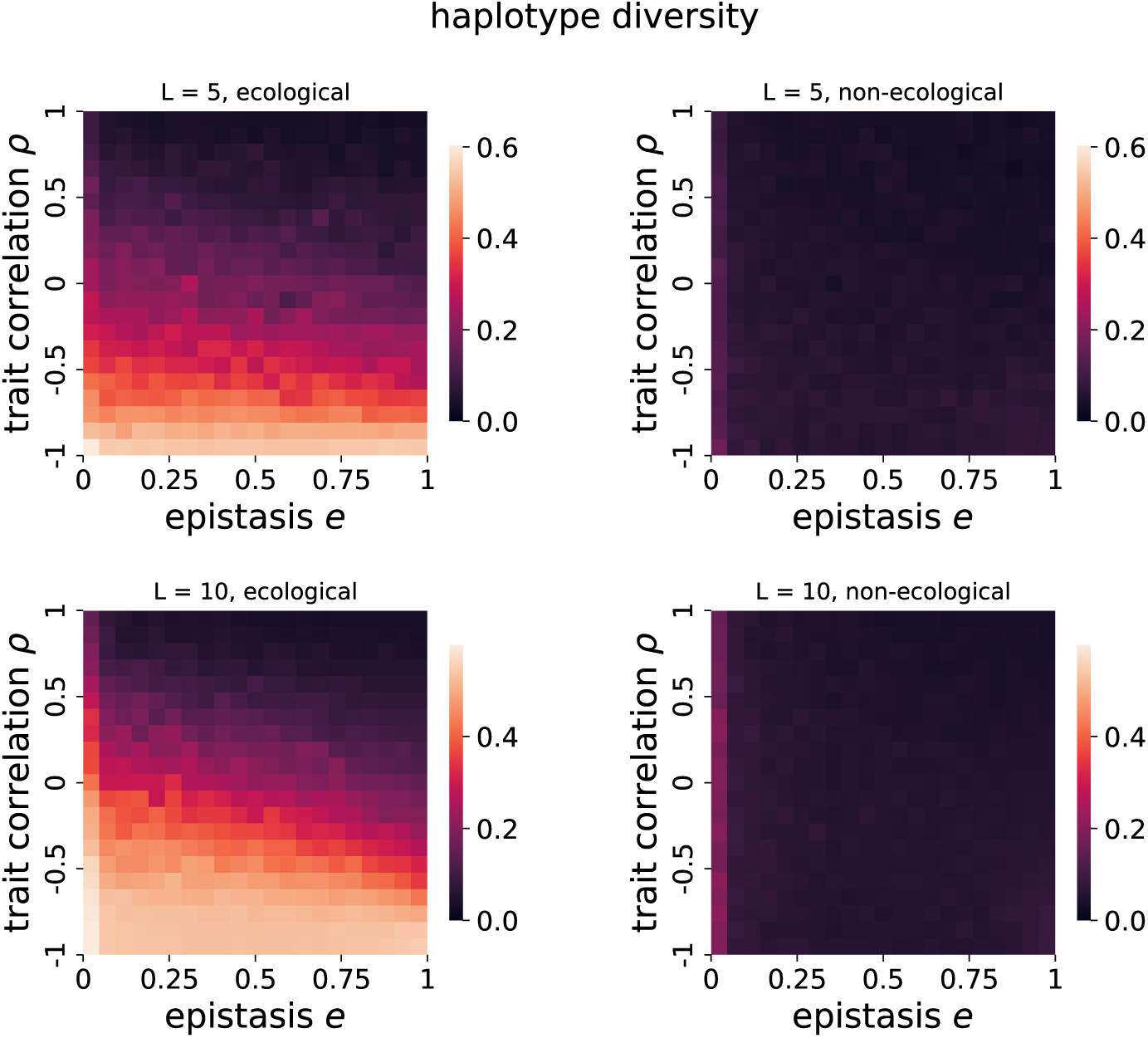
Haplotype diversity tends to decrease with increasing strength of epistasis. The graphs show the mean haplotype diversity *h* at the final state for the ecological and the non-ecological model. The top row shows the results for *L* = 5 loci, and the bottom row for *L* = 10 loci. The left column shows the ecological model, and the right column shows the non-ecological model. Each point represents an average obtained from 100 different fitness landscapes with the same parameters with 20 replicates per landscape. The remaining parameters are the population size *N* = 1000, the mutation rate per locus *µ* = 0.001, the amount of resources *s_r_* = 1, and the variance of fitness effects *σ* = 0.1.

**Figure S4:**
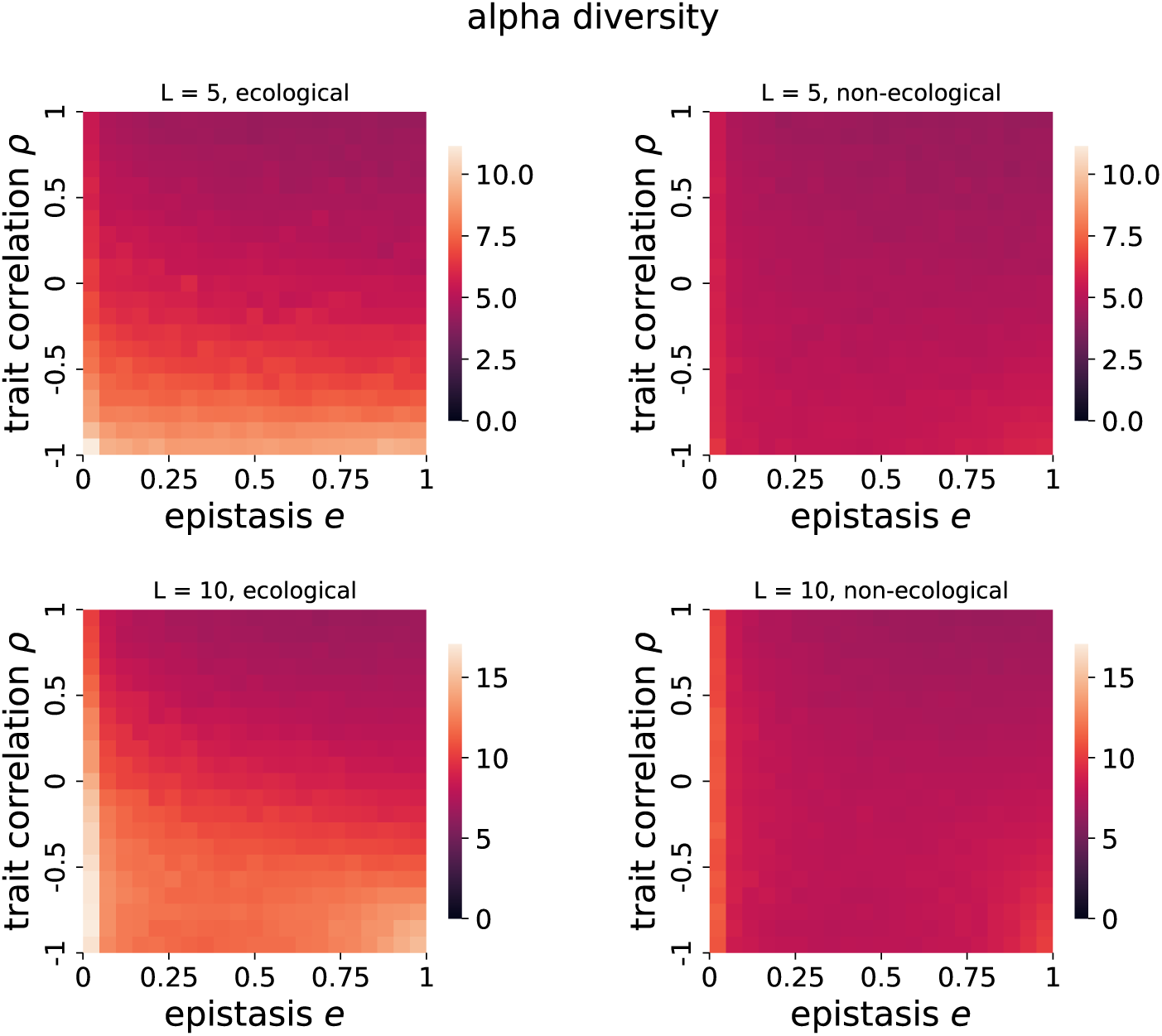
Alpha diversity. The graphs show the mean alpha diversity at the final state for the ecological and the non-ecological models. The top row shows the results for *L* = 5 loci, and the bottom row for *L* = 10 loci. The left column shows the ecological model, and the right column shows the non-ecological model. Each point represents an average obtained from 100 different fitness landscapes with the same parameters with 20 replicates per landscape. The remaining parameters are the population size *N* = 1000, the mutation rate per locus *µ* = 0.001, the amount of resources *s_r_* = 1, and the variance of fitness effects *σ* = 0.1.

